# A case of mycobacteriosis associated with *Mycobacterium pseudoshottsii* in aquarium-reared fish in Japan

**DOI:** 10.1101/2022.07.14.500145

**Authors:** Takeshi Komine, Ihara Hyogo, Ono Kentaro, Mitsumi Yoshida, Yuma Sugimoto, Mari Inohana, Hanako Fukano, Osamu Kurata, Shinpei Wada

## Abstract

In 2019, several aquarium-reared fish died at a sea life park in Japan. Necropsy revealed micronodules on the spleen in the dotted gizzard shad (*Konosirus punctatus*). Seven of 16 fish exhibited microscopic multifocal granulomas associated with acid-fast bacilli in the spleen, kidney, liver, alimentary tract, mesentery, gills, and/or heart. Bacterial cultures yielded isolates from the dotted gizzard shad and a Japanese sardine (*Sardinops melanostictus*). Microbiological examination, multilocus sequence typing analysis, and variable number of tandem repeats analysis with a newly proposed 6-loci set revealed the isolates as *Mycobacterium pseudoshottsii*. To our knowledge, this is the first isolation of *M. pseudoshottsii* from aquarium-reared fish.

---

*Mycobacterium pseudoshottsii*, a slow-growing, photochromogenic, nontuberculous mycobacterium, is a member of the *Mycobacterium marinum* group (MMG; *M. marinum, M. ulcerans, M. pseudoshottsii, M. shottsii*, and *M. liflandii*), whose members are closely related genetically [10, 22, 24]. MMG species are mycolactone-producing mycobacteria, and mycolactone is a cytotoxic macrolide toxin whose gene is encoded on pMUM plasmids [9, 21].

*M. pseudoshottsii* is reported to have caused infectious diseases in at least ten species of fish. In Chesapeake Bay, the pathogen was initially isolated from wild striped bass (*Morone saxatilis*) in 2005 [22] and from wild white perch (*Morone americana*) in 2007 [25]. Nakanaga et al. isolated the bacteria from yellow tail (*Seriola quinqueradiata*), greater amberjack (*Seriola dumerili*), striped jack (*Pseudocaranx dentex*), sevenband grouper (*Epinephelus septemfasciatus*), and yellowtail amberjack (*Seriola lalandi*) farmed in western Japan [18]. Mugetti et al. reported *M. pseudoshottsii*-associated mycobacteriosis in red drum (*Sciaenops ocellatus*), European sea bass (*Dicentrarchus labrax*), and gilthead sea bream (*Sparus aurata*) on farms in the Mediterranean Sea [17].

Variable number of tandem repeats (VNTR) analysis is a valuable molecular typing method based on copy numbers of 40–100-bp repetitive sequences in the genome [29]. VNTR analysis is sometimes applied to subdivide mycobacterial species with high genetic homology, as observed among MMG species [6, 27, 31].

Tokyo Sea Life Park, an aquarium in Japan, rears and exhibits approximately 50 fish from over ten species in a single tank, including the Japanese sardine (*Sardinops melanostictus*), dotted gizzard shad (*Konosirus punctatus*), marbled sole (*Pleuronectes yokohamae*), bluefin searobin (*Chelidonichthys spinosus*), horse mackerel (*Trachurus japonicus*), Japanese butterfish (*Psenopsis anomala*), Japanese conger (*Conger myriaster*), Japanese seabass (*Lateolabrax japonicus*), Japanese whiting (*Sillago japonica*), and white croaker (*Argyrosomus argentatus*). The fish were captured in Tokyo Bay and housed in an elliptical/cylindric tank with an approximately 9-t capacity, closed-circulation system and water temperature of approximately 20°C. Several fish reared in this tank have died each month since February 2019 from nontuberculous mycobacteria. A field test (AFB-Color staining kit, Merck KGaA, Darmstadt, Germany) conducted on a few dead fish revealed acid-fast bacilli (AFB). Therefore, we performed bacterial cultures and molecular biological examinations to isolate and identify the AFB.

We collected six freshly dead fish and ten live fish from the aforementioned tank. The live fish were euthanized via cervical transection after anesthetization with an overdose of 2-phenoxyethanol in accordance with the American Veterinary Medical Association Guidelines for the Euthanasia of Animals (2013 edition) [15]. The collected fish were frozen at −20°C until necropsy. The fish were thawed and routinely dissected, and their external and internal gross features were observed. Tissues were collected from the major organs (i.e., spleen, liver, kidney, heart, gills, and/or alimentary tract), fixed in 10% phosphate-buffered formalin solution, routinely processed and embedded into paraffin blocks for histopathology. For the spleens, livers, and kidneys, parts of the tissues were frozen for microbiological and molecular biological analyses. The tissue blocks were sectioned and stained with hematoxylin and eosin, Ziehl-Neelsen stain, and a modified Nyka’s method as described by Harada (1977) [11, 20]. Mycobacteria were isolated as follows. Frozen tissue samples from 16 fish were homogenized with 0.5 ml of phosphate-buffered saline (PBS) (-) and decontaminated with 1 ml of N-acetyl-L-cysteine-sodium citrate (NALC)-NaOH or 0.75 ml of 1N HCl. After decontamination for no more than 15 min, the NALC-NaOH-treated samples were neutralized by adding at least 6 ml of 0.067 M phosphate buffer adjusted to pH 6.8, and the HCl-treated samples were neutralized by adding 0.75 ml of 1N NaOH. The samples were centrifuged at 3,000 ×g for 20 min, then the supernatant was discarded, and the pellet was resuspended in 1 ml of Middlebrook 7H9 broth base (Becton, Dickinson and Company, Franklin Lakes, NJ, USA) supplemented with 10% BBL Middlebrook Oleic Albumin Dextrose Catalase (OADC) enrichment (Becton, Dickinson and Company). Aliquots (25 μl of each sample) were inoculated on Middlebrook 7H10 agar supplemented with 10% OADC enrichment and spread on 2% Ogawa egg slant (Kyokuto Pharmaceutical Industrial Co., Ltd., Tokyo, Japan). The media were incubated at 25°C for 2 months and were checked daily for the first week, then once weekly thereafter. A single colony was subcultured on Middlebrook 7H10 agar to obtain pure isolates. The isolates were stained with Ziehl-Neelsen on glass slides to check their acid-fast stainability. Two of the three mycobacterial isolates obtained from a dead fish and a live fish (named NJB1907-Z4 and NJB1907-f19, respectively; Supplemental Table 1) were subjected to the following examinations.

To classify the two isolates based on the Runyon classification system [23], their pigment production ability was tested on 2% Ogawa egg slants as per Fukano et al. (2015) [7]. The characteristics of the isolates and type strains (*M. marinum* JCM 17638 and *M. pseudoshottsii* JCM 15466), such as arylsulfatase and Tween 80 hydrolysis, were tested according to the procedure described by the Japanese Society for Tuberculosis (2016) [30] with a modification of the incubation temperature to 25°C. The arylsulfatase test was conducted on days 3 and 14 [8].

Genomic DNA was extracted from five strains (*M. marinum* JCM 17638, ATCC BAA535, *M. pseudoshottsii* JCM 15466, NJB1907-Z4, and NJB1907-f19) following the method of Komine et al. (2021) [13] and subjected to the subsequent experiments. Multilocus sequence typing (MLST) analysis was conducted based on a total of 1,997 bp of concatenated sequences of three housekeeping genes: 913 bp of 16S rRNA, 683 bp of RNA polymerase b-subunit (*rpoB*), and 401 bp of 65-kDa heat-shock protein (*hsp65*) following the method of Komine et al. (2021) [13]. The obtained sequences for the three genes from NJB1907-Z4 were deposited into the DNA Data Bank of Japan under accession numbers LC699671, LC699669, and LC699670, respectively. Presence of the insertion sequences, *IS2404* and *IS2606*, relating to mycolactone production in the isolates was confirmed using the primer sets MU5–MU6 and MU7–MU8 through the cycles as described by Stinear et al. (1999) [26].

VNTR analysis was conducted for six loci: MIRU5, MIRU9 [28], loci 4, 6, 8, and 16 [1], and a new locus V4. Supplemental Table 2 shows the primer sets used. The V4 region was extracted from the draft genome of *M. marinum* NJB1728-216S (BQLD01000000) using Tandem repeats finder (TRF) version 4.09 (https://tandem.bu.edu/trf/trf.html) [3]. The primer set of V4F (5’-ATCCAGTCAGGAATGTCGGC-3’) and V4R (5’-AGGCCAAGTGGTTCTGGTTC-3’) was designed in the flanking region of V4 with Primer-BLAST (https://www.ncbi.nlm.nih.gov/tools/primer-blast/).

VNTR profiles were determined using three methods as follows. In the first method, estimated sequences of six loci from eight strains, *M. marinum* ATCC BAA535 (CP000854.1), MMA1 (CP058277.1), JCM 17638 (AP018496.1), *M. pseudoshottsii* JCM 15466, *M. liflandii* 128FXT (CP003899.1), *M. shottsii* JCM 12657 (AP022572.1), *M. ulcerans* subsp. *shinshuense* ATCC 33728 (AP017624.1), and *M. ulcerans* SGL03 (LR135168.1), were obtained via *in silico* polymerase chain reaction (PCR) using BLAST (https://blast.ncbi.nlm.nih.gov/Blast.cgi). From the sequences, repeat numbers were determined using TRF. If no repeat numbers were found with TRF, they were determined from alignment between the sequences and repeat sequences obtained from other strains with TRF, using ClustalW in MEGA X [14]. In the second method, *in vitro* (laboratory-observed) PCR was conducted on three strains (*M. marinum* JCM 17638, ATCC BAA535, and *M. pseudoshottsii* JCM 15466) using the above-listed primer sets. Loci 6, 8, 16 and V4 were amplified under the cycle conditions of Ablordey et al. (2005) [1]. MIRU 5 and 9 were amplified under the cycle conditions of Stragier et al. (2005) [28]. PCR was performed in a total of 20 μl of PCR mixture containing 11.3 μl H_2_O, 4.0 μl 5× colorless buffer, 1.6 μl 2.5 mM dNTP, 1.0 μl each of 10-μM forward and 10-μM reverse primers, 0.1 μl GoTaq^®^ DNA polymerase (Promega Corp., Madison, WI, USA), and 1 μl of template DNA. The amplicons were purified using NucleoSpin^®^ Gel and PCR Clean-up (MACHEREYNAGEL GmbH & Co. KG, Düren, Germany). The purified amplicons were sequenced by FASMAC (Kanagawa, Japan). Repeat numbers were determined from the sequences using TRF and MEGA X. In the third method, VNTR profiles of NJB1907-Z4 and NJB1907-f19 were determined by comparing the two strains and three strains (used in the second method) in gel electrophoresis. PCR was conducted as described above. Amplicons were detected by gel electrophoresis in 2% agarose-1 ×Tris-borate-EDTA buffer at 100 V for 40 min. From these VNTR profiles, a minimum spanning tree was generated with PHYLOViZ (https://online.phyloviz.net/index).

Gross examination revealed a few white micronodules of approximately 0.5 mm in diameter on the spleens of two dotted gizzard shad (1907-f16, f19), skin ulceration in one fish (1907-f7), and exophthalmos in two fish (1907-f5, f14). Microscopic examination revealed AFB-associated multifocal granulomas in the serosal membrane to muscular layers of the alimentary tract, mesentery, interstitial kidney tissue, and/or parenchyma of the liver and spleen in a Japanese sardine, four dotted gizzard shad, and the marbled sole (Supplemental Table 1). The granulomas often had a necrotic center with AFB, surrounded by epithelioid cells and a thin outermost rim of fibroblasts and connective tissue (Fig. 1). Some fish exhibited granulomas composed of mononuclear macrophages without epithelioid cells and a thin outermost rim.

**Fig. 1.**
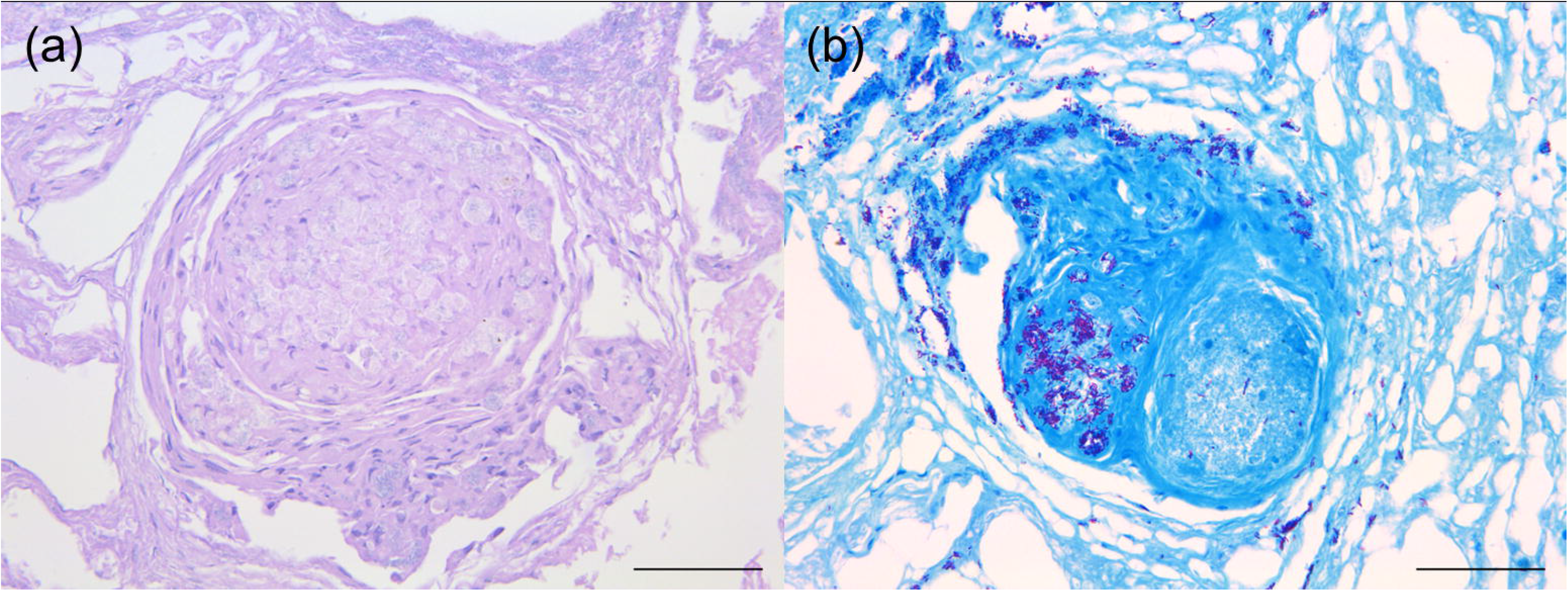
Epithelioid granulomas in the mesentery of an infected fish (ID 1907-f16). A necrotic area with acid-fast bacilli, surrounded by epithelioid cells and a thin outermost rim of fibroblasts and connective tissue was observed. (a) hematoxylin and eosin stain; (b) Ziehl-Neelsen stain. Bars = 50 μm.

Runyon classification, biochemical examination, and insertion sequence detection results indicated that isolates NJB1907-Z4 and NJB1907-f19 showed similar characteristics to those of *M. pseudoshottsii* JCM 15466 (Table 1). The isolates were also closely related to *M. pseudoshottsii* JCM 15466 on the concatenated and minimum spanning trees (Figs. 2, 3) as per their VNTR profiles (Supplemental Table 3). These results suggested that isolates NJB1907-Z4 and NJB1907-f19 were *M. pseudoshottsii. M. pseudoshottsii* has been isolated from several wild and cultured fish. However, to our knowledge, no infectious case of the pathogen has been previously reported in aquarium-reared fish; this is the first isolation of *M. pseudoshottsii* from aquarium-reared fish.

**Table 1.**
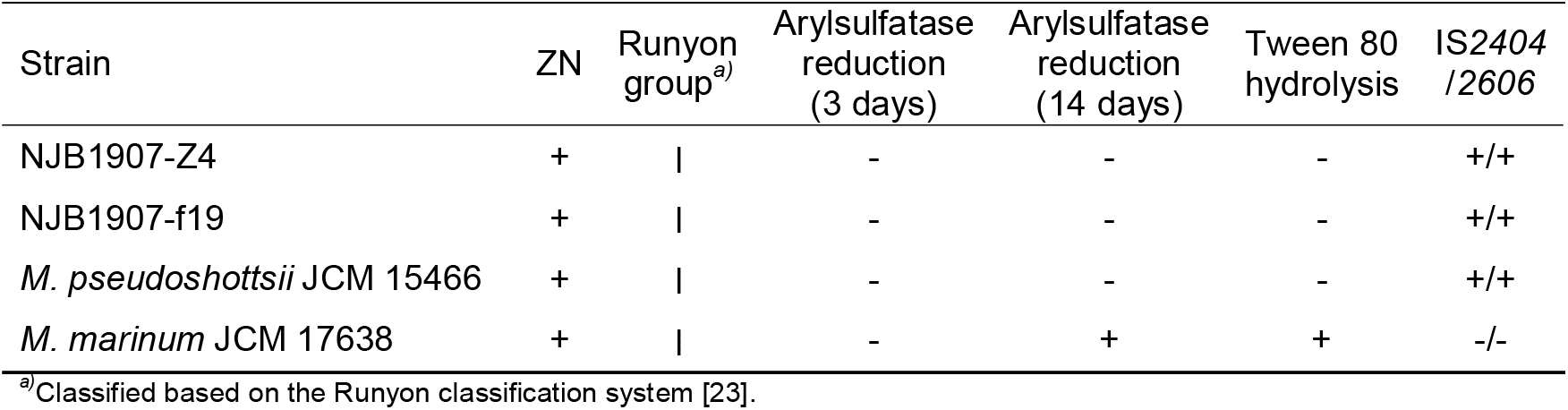
Phenotypic and genomic characteristics of the isolates and type strains

**Fig. 2.**
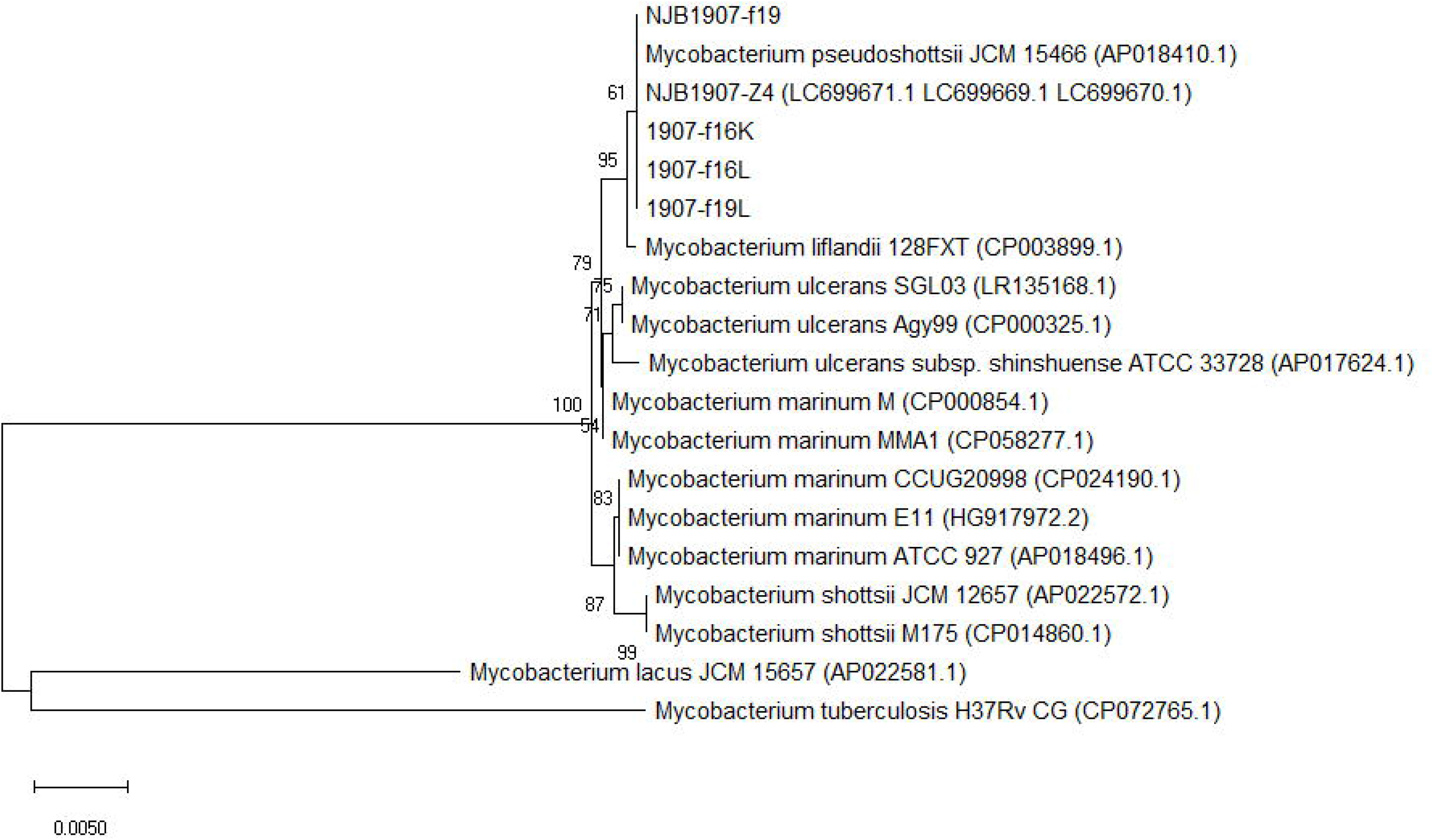
Phylogenetic tree generated from a concatenated 1,997-bp sequence of the 16S rRNA, *rpoB*, and *hsp65* genes from a kidney (1907-f16K), livers (1907-f16L, f19L) and two isolates (NJB1907-Z4, f19). The tree was constructed using the neighbor-joining method with Kimura’s two-parameter model in Mega X [14]. Bootstrap values are shown at nodes as percentages of 1,000 replicates. GenBank accession numbers are in parentheses.

**Fig. 3.**
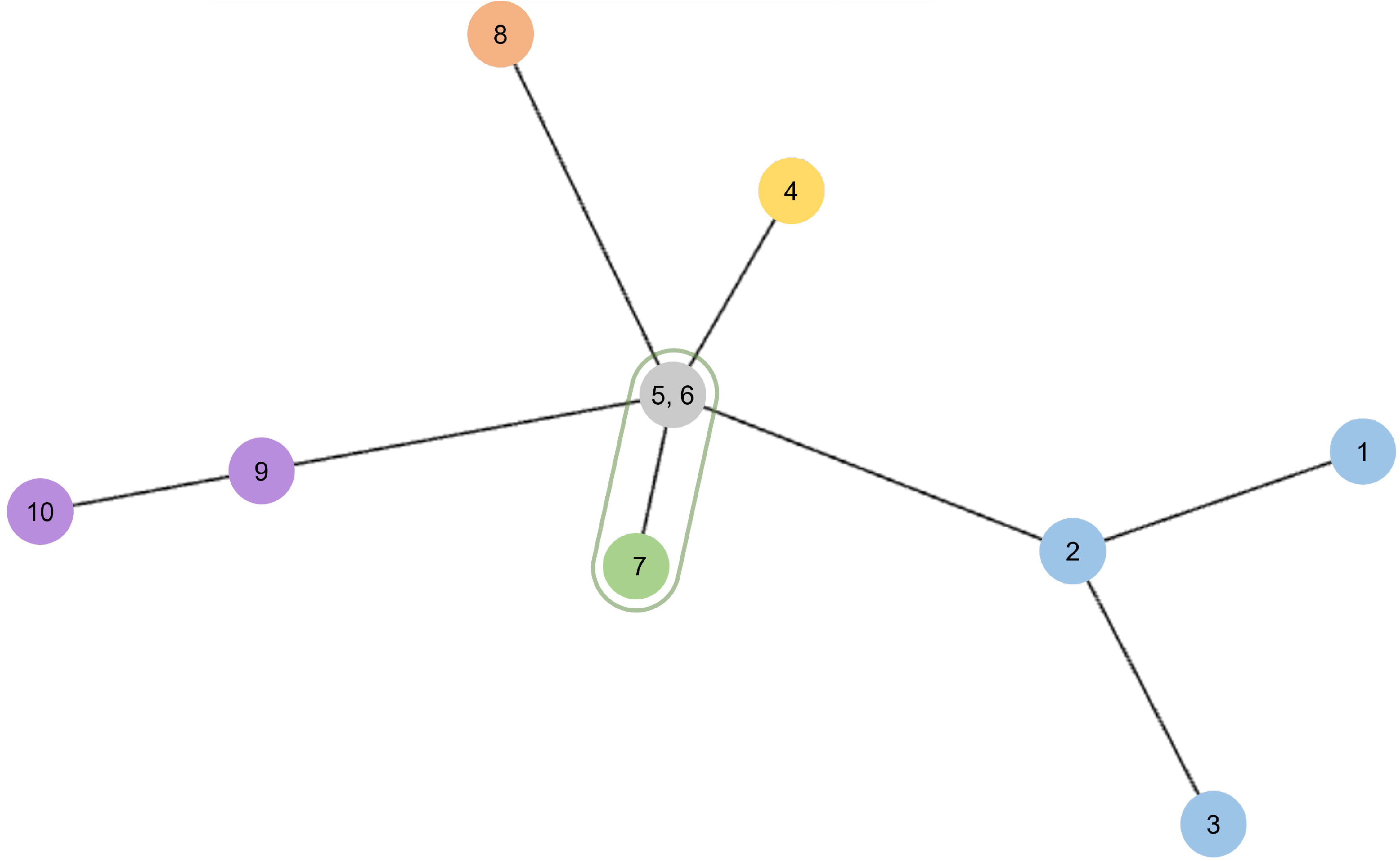
Minimum spanning tree based on variable number of tandem repeats profiles from the isolates and type strains. The tree was generated using the goeBURST algorithm in PHYLOViZ software (https://online.phyloviz.net/index). 1. *Mycobacterium marinum* ATCC BAA535; 2, *M. marinum* MMA1; 3, *M. marinum* JCM 17638; 4, *M. liflandii* 128FXT; 5, NJB1907-Z4; 6, NJB1907-f19; 7, *M. pseudoshottsii* JCM 15466; 8, *M. shottsii* JCM 12657; 9, *M. ulcerans* subsp. *shinshuense* ATCC 33728; 10, *M. ulcerans* SGL03.

Because MMG species are reported to be highly genetically homologous, they are difficult to identify based only on sequencing analysis of the 16S rRNA gene [4]. Therefore, they should be identified using adequate methods as described herein, including biochemical examinations (arylsulfatase, Tween 80 hydrolysis), MLST analysis based on the housekeeping genes (16S rRNA, *rpoB, hsp65*), and VNTR analysis of MIRU5 and 9 and loci 4, 6, 8, 16, and V4. The present study revealed that the biological, biochemical, and molecular biological examinations used can help subdivide MMG species in greater detail.

*M. marinum*, a representative MMG species, is a common causative agent of mycobacteriosis in fish and causes a skin infection known as “fish tank granuloma” in humans, thus posing a zoonotic concern [12]. Regarding the effects on aquatic resources, human economy and public health, appropriate measures, including sanitation, disinfection of the facility, and eradication of carrier fish, are necessary as primary control strategies to control *M. marinum* infection in fish [2, 5, 16, 19]. However, *M. pseudoshottsii* has never been isolated from humans. Thus, the risk of *M. pseudoshottsii* infection in humans may be lower than that of *M. marinum* infection. In conclusion, the pathogenesis of each clinical case should be clarified, and the causative agent should be identified in detail using adequate methods as described herein to avoid applying the wrong countermeasures to suspected clinical cases associated with MMG.

## Potential conflicts of interest

None declared.

## Acknowledgment

We thank Traci Raley, MS, ELS, from Edanz (https://jp.edanz.com/ac) for editing a draft of this manuscript.

## Notes

### Competing Interest Statement

The authors have declared no competing interest.

